# Ancestral Protein Reconstruction of a membrane trafficking GTPase uncovers unanticipated properties of the ancestral protein and of modern Arf1 GTPases

**DOI:** 10.1101/2025.10.27.684887

**Authors:** Marine Alves, Mandeep Sivia, Kristína Záhonová, Catherine L. Jackson, Joel B. Dacks

**Author notes:** to whom correspondence should be addressed CLJ and JBD.

## Abstract

The emergence of eukaryotes from their prokaryotic ancestors (eukaryogenesis) marked a fundamental shift in cellular organisation, with the appearance of intracellular compartments including the nucleus, the Golgi apparatus and endosomes. These organelles are part of the endomembrane system of eukaryotic cells, which mediates many processes, including secretion of proteins to the exterior of the cell, uptake of material by endocytosis, and compartmentalized degradation of cellular components. The period of eukaryogenesis after the merger of prokaryotic lineages but preceding the last eukaryotic common ancestor, is inferred to have involved a progressive increase in cellular complexity through expansion of organelle-specific protein machineries. However, the steps and stages of organelle emergence during this period are poorly understood as no extant organisms exist from this period, precluding the use of comparative genomics to determine the properties of ancestral proteins present. Membrane trafficking pathways linking organelles are regulated by Arf family GTPases, including Arf1 and Arf6, both present in the last eukaryotic common ancestor. Here we use ancestral sequence reconstruction and molecular cell biological characterization to explore the properties of the ancestor of the Arf1 and Arf6 GTPases. Arf1 has a major function at the Golgi apparatus in regulation of the secretory pathway, whereas Arf6 regulates endocytic pathways at the plasma membrane and endosomes. Our results indicate that the ancestral Arf1/6 protein localizes to both the Golgi and the plasma membrane. We find that localization to the plasma membrane is due to a C-terminal polybasic motif that unexpectedly is also found in a number of modern Arf1 proteins from a wide diversity of eukaryotes. Our data suggest that the ancestral Arf protein acted at both internal compartments and the cell periphery, a feature preserved in a number of modern Arf1 proteins.

## Introduction

The evolution of eukaryotes from their prokaryotic ancestors was an important transition in the history of life, resulting in nearly all complex life forms on earth today. Understanding of this series of events, known as eukaryogenesis, has been the focus of considerable research over many years and has seen significant progress recently, but many questions remain unanswered ^1^. Eukaryotes arose from a symbiosis between at least two prokaryotic lineages. These lineages include an alphaproteobacterium that gave rise to the mitochondria, and an archaeal cell, that contributed at least some of the other organellar machinery ^1–3^. The resulting endosymbiont then evolved, acquiring hallmark eukaryotic traits and culminating in a population of single-celled eukaryotic organisms, the Last Eukaryotic Common Ancestor (LECA), which gave rise to the extant lineages inhabiting Earth today ^1^. The most powerful and frequently used approach to understanding eukaryogenesis through these ancestors is comparative genomics of extant descendant organisms. For eukaryotic origins, the challenging investigation of the contributing prokaryotic lineages and earliest eukaryogenic events has advanced rapidly, with recent studies describing the Asgardarchaeota, from which the eukaryotic host is derived ^2^, as well as the study of the closest alphaproteobacterial relatives of mitochondria ^4^. The LECA marks the end of the eukaryogenic period and is a highly tractable reconstruction point by comparative genomics ^5^. From a wide range of analyses, it is inferred to have possessed nearly the complete set of organelles observed in many modern cells. This configuration is underpinned by a large genome complement of molecular machinery, characterized by paralogous expansions of protein families ^5^. Though challenges remain, this LECA reconstruction is considered to be robust and comprehensive.

One of the major hallmarks of eukaryotic cells that distinguish them from their prokaryotic ancestors is the endomembrane system^1^. This interconnected system of internal membrane-bound organelles, including the endoplasmic reticulum (ER), the Golgi apparatus, endosomes and lysosomes, allowed compartmentalization of biochemical activities and presumably provided a tremendous evolutionary advantage. The secretory pathway assures the delivery of secreted proteins, such as signalling molecules or proteins for predation and defence to the exterior of the cell, and membrane proteins to intracellular locations. The endocytic pathway assures the uptake of extracellular material including nutrients, as well as internalization of plasma membrane (PM) proteins such as signalling receptors for recycling or transport to the lysosome for degradation. In both the secretory and endocytic pathways, communication between organelles is assured by membrane-bound vesicular carriers^6^. Thus, these pathways not only ensured specificity in secretion and internalization but also provided an interconnecting network between many organelles, allowing localized concentration of proteins and lipids for various functions.

The archaeal ancestor of eukaryotes provided some of the precursors of the protein machinery for vesicular transport found in extant eukaryotes ^7–9^. Between this first eukaryotic ancestor and the LECA, extensive elaboration of the endomembrane system occurred, producing the endomembrane organelles. The LECA is inferred as having possessed a highly sophisticated set of membrane trafficking machinery, much of which is comprised of protein families, where the duplicated family members carry out similar biochemical functions, but at different transport pathways or organelles ^5,10^. Thus a full understanding of eukaryogenesis requires explanation of how the endomembrane system evolved.

The molecular machinery of membrane traffic in the endomembrane system includes regulators such as the Arf family GTPases, which are universally present in eukaryotes ^11,12^. The Arf family includes Arf, Arf-like (Arl) and Sar GTPases, which are involved in different steps of membrane traffic including vesicle budding, vesicle tethering, organelle movement, cytoskeleton-membrane interactions and lipid homeostatsis ^13–15^. The Arf family GTPases carry out their functions by cycling between GDP- and GTP-bound forms, where the GTP-bound form is tightly membrane bound and recruits effector proteins to the membrane. Among small GTPases, Arf family proteins are unique in having an N-terminal amphipathic helix, often N-myristoylated, that promotes membrane binding of the active GTP-bound form ^16^. Arf1 and Arf6 are hence soluble in their GDP-bound form, and the change in conformation of the N-terminal AH upon GTP binding promotes tight membrane association of the active form. In cells, the level of the soluble pool of an Arf protein depends on its level of activation and also on the presence of membrane-associated receptors for the GDP-bound form ^17–24^.

The Arf family is ancient, as the eukaryotic proteins all arose from an ancestral group, the ArfR GTPases, which were present in the archaeal ancestor of eukaryotes ^8,25^. The LECA is inferred to have already possessed 15 members of the Arf GTPase family ^12^. The originally identified Arf1-6 proteins in mammalian cells, and their orthologues throughout eukaryotes, form a single group within the Arf family. LECA is predicted to have had both Arf1 and Arf6 proteins ^12^. Arf1 has expanded in many lineages, including in mammals to produce Arf1-5 ^16^. Arf1 carries out essential functions in the secretory pathway, including vesicle formation at the Golgi, while Arf6 is involved in endocytosis pathways and actin cytoskeleton regulation at the PM ^13,14,26,27^. Arf1 is one of two Arf family proteins found in every eukaryote examined to date, and Arf1 proteins are highly conserved at the primary sequence level. For example, human Arf1 is 88% identical to the protist *Planomonas micra* Arf1a, and 77% identical to the yeast *Saccaromyces cerevisiae* Arf1. Arf6, on the other hand, has been lost in a number of eukaryotic lineages, including early in the evolution of the Discoba and of the Archaeplastida (land plants and red, green and glaucophytes algal relatives), and Arf6 sequences are more divergent at the sequence level ^12^. For example, human Arf6 is only 51% and 60% identical to the Arf6 proteins of *Planomonas micra* and *Saccaromyces cerevisiae*, respectively. Because Arf1 and Arf6 are each other’s closest relatives within the Arf Family ^12^, their emergence likely represents a relatively late step in eukaryogenesis, close to the LECA.

Comparative genomic analyses based on existing eukaryotic genomes can infer the predicted proteins present in LECA. When Asgard archaeal genomes are also included in comparative genomic analyses, this approach can predict the genes and their protein products that were present in the common ancestor of eukaryotes and the Asgard archaea, which predates LECA and the mitochondrial symbiosis event. However, this approach cannot be used to predict the features of our ancestors between these two points in the evolutionary time line, as no extant organisms originating from this evolutionary period have been identified. Given the complexity and sophistication of the LECA, this period is crucial to our understanding of eukaryogenesis, and hence new approaches are required to provide advances in this area. Because the LECA is inferred to have possessed many paralogues of membrane trafficking machinery ^10^*inter alia*, the challenge is to reconstruct the duplication steps giving rise to that complement. Determining the properties of the pre-duplicated ancestral proteins is precisely what is needed to reconstruct the most recent steps to LECA.

Ancestral Protein Reconstruction (APR) allows for the prediction of ancestral protein sequences (i.e. Ancestral Sequence Reconstruction, ASR) and then the experimental testing of the biochemical and/or cell biological properties of the predicted proteins^28,29^. This approach has been used successfully for metabolic enzymes, transcription factors and structural proteins and back to deep evolutionary time points, including LECA proteins ^28,30,31^. In this study, we have used APR to move beyond the LECA boundary to reconstruct the ancestor of Arf1 and Arf6 GTPases and to explore whether the cell biological properties of the predicted Arf1/6 ancestral sequences resemble either Arf1, Arf6 or a combination of both.

## Results

### Ancestral Sequence Reconstruction

In order to perform ASR of the ancestor of Arf1 and Arf6, it was first necessary to construct a broadly sampled and representative dataset. Starting with the dataset from Vargová et al. 2021 ^12^, we updated the taxon sampling using genomes or transcriptomes of key eukaryotic lineages that had been sequenced in the intervening period and removed organisms redundant to our overall aim of a broad and even sampling of eukaryotes (Supplementary Table 1). Because the aim of the project was reconstruction of nodes in the Arf tree pre-LECA, much more recent, lineage-specific, expansions and highly divergent sequences were identified by preliminary phylogenetic analyses (Supplementary Figure 1A-D), and removed from the dataset. These analyses were carried out for Arf1, Arf6, as well as their closest sister groups Arl1 and Arl5. The latter two subfamilies form a clade that is the closest to the Arf1 - Arf6 clade in the Vargova et al. 2021^12^ analysis. As such we include them as outgroups to polarize the analysis under the assumption that the rooting of the entire Arf family does not lie within this set of sequences. The least divergent representatives of each subfamily were aligned and taxon-specific insertions were masked out such that they were not included in the ancestrally reconstructed sequence. This resulted in a final dataset of 363 sequences.

Under the assumption that the genes in question are vertically inherited ^12^, a constrained tree was constructed following the species relationships of the taxa sampled ^32,33^ (Supplementary Figure 2), with an assumed rooting on the mid-point branch between the Arf1+Arf6 and the Arl1+Arl 5 clades (see Methods). The branch lengths in the Arf6 clade are substantially longer than in the Arf1 clade, consistent with past observations of negative selection on Arf1 sequence evolution and divergence in Arf6 (Figure 1, Supplementary Figure 2). ASR was undertaken using this guide tree in both PAML and IQ-TREE (Figure 2A). Initial reconstruction was performed with the optimally chosen model (LG+R8) by the embedded ModelFinder in IQ-TREE. In order to test the effect of a mixture model, we implemented a (LG+C60+R10) model within the IQ-TREE which was determined to be the optimal model and used for final ASR (Supplementary Table 2).

**Figure 1:**
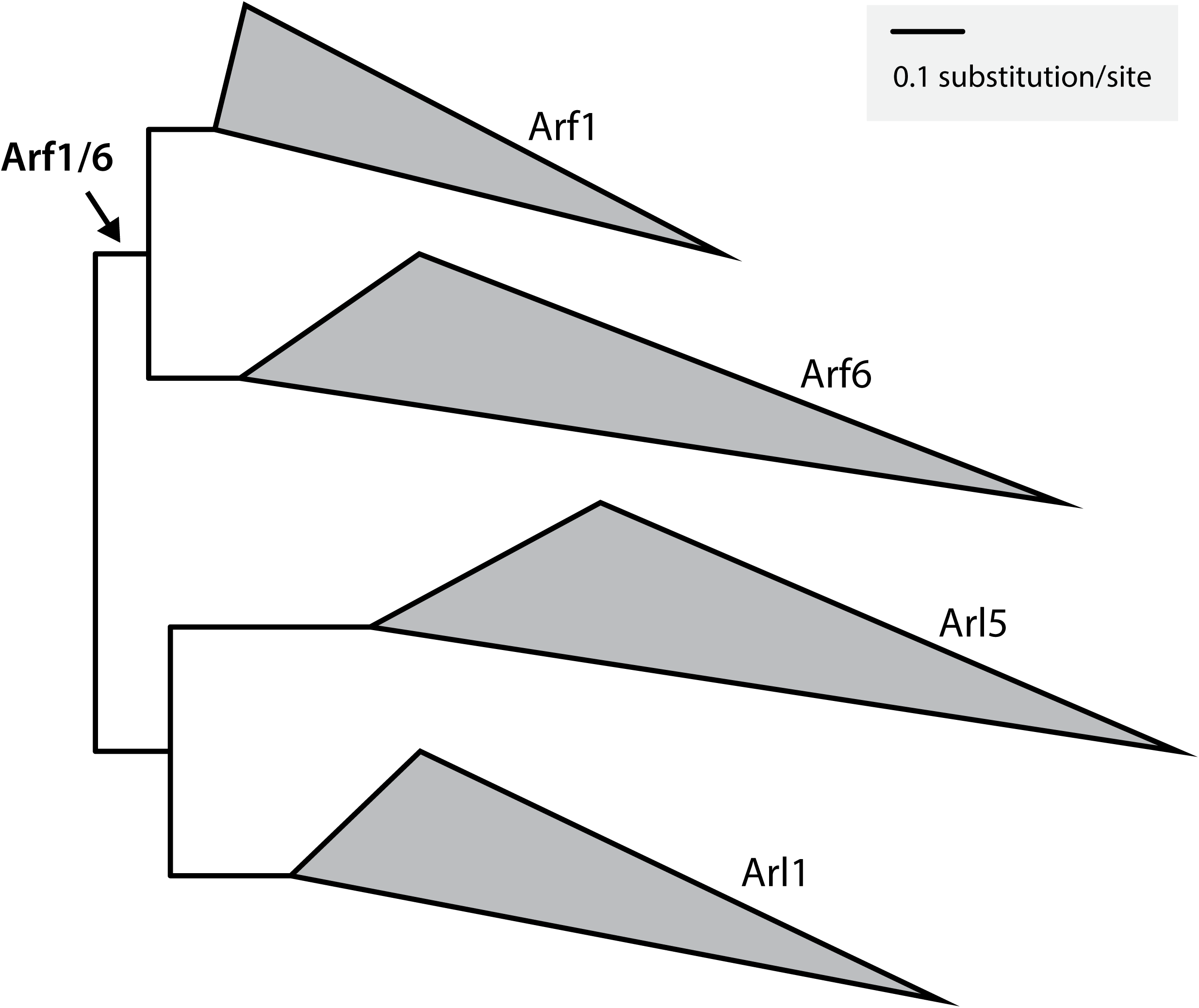
Illustrated constraint tree calculated for Arf1, Arf6, Arl1, and Arl5. This phylogeny shows the more divergent nature of Arf6 sequences as compared with Arf1. Clades are collapsed for clarity (for full tree see Supplementary Figure 2).

**Figure 2:**
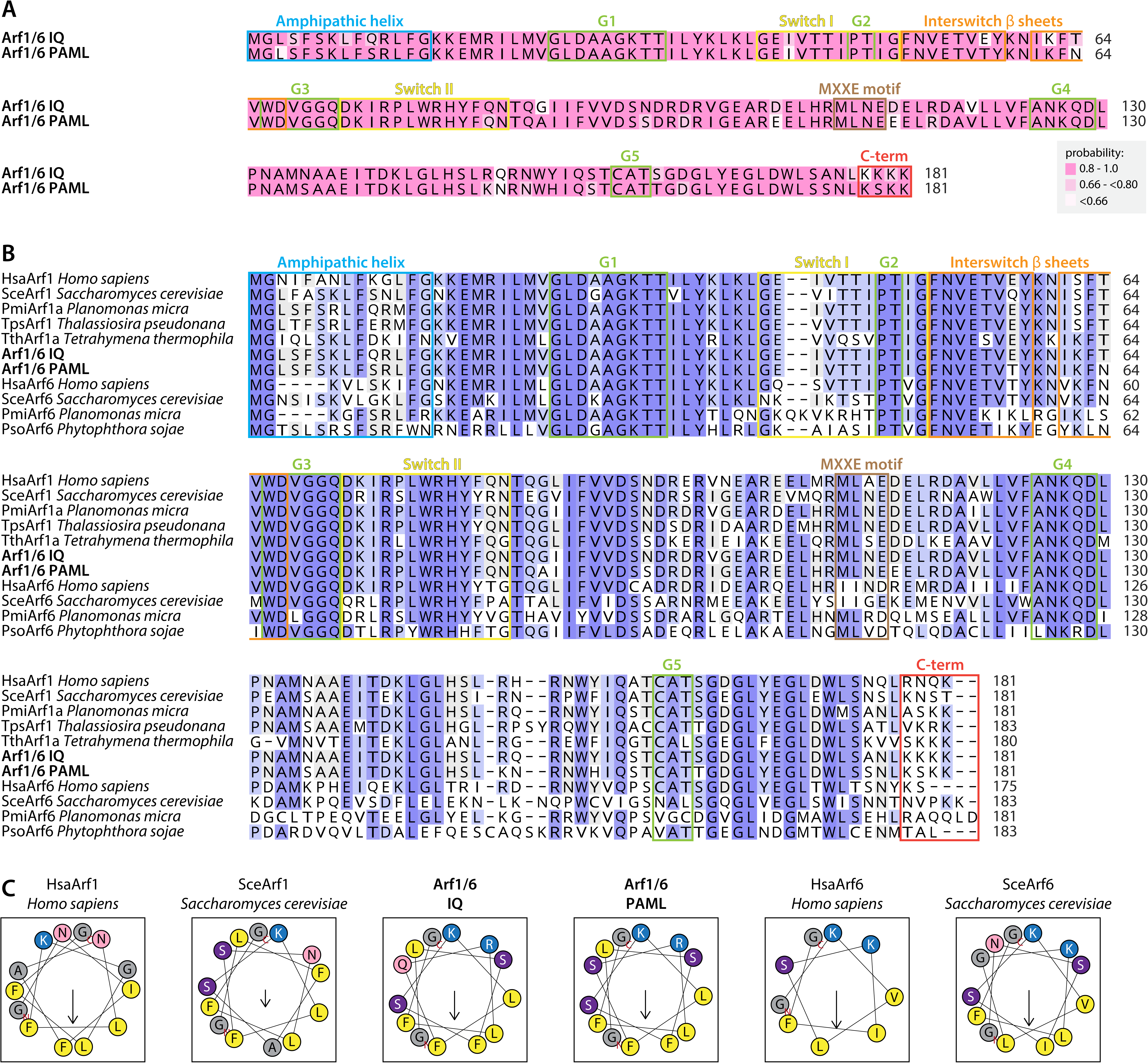
Properties of the Arf1/6 ancestral proteins predicted by PAML and IQ-TREE. (A) Alignment of the two predicted ancestral sequences, Arf1/6PAML and Arf1/6IQ. Key G-Boxes and and sequence motifs are annotated above. Residue-specific probabilities are colour-coded as inset. (B) Alignment of Arf1/6PAML and Arf1/6IQ predicted ancestral sequences and selected eukaryotic Arf1 and Arf6 proteins. Identical residues in 80% or more (all but 2) of the sequences are indicated by dark purple boxes; in 70% (all but 3) by light purple boxes. Same motifs as in part A are annotated. (C) Helical wheel plots of N-terminal AHs of some of the sequences shown in part B. Helical wheel representations of the N-terminal AHs, generated by Heliquest ^67^, are shown. Color code for the amino acids is yellow: Val, Leu, Ile, Met, Phe, and Trp; dark purple: Ser and Thr; red: Asp and Glu; pink: Asn and Gln; blue: Lys and Arg; grey: other residues. Arrow represents the helical hydrophobic moment, with length proportional to the calculated value and direction indicating that the helix is amphipathic in the perpendicular direction.

Our ancestral reconstructions focused on three nodes of interest, the ancestor of all Arf1s, the ancestor of all Arf6s, and in particular the Arf1/6 ancestor. The ancestral Arf1/6 sequences predicted by PAML (Arf1/6PAML) and IQ-TREE (Arf1/6IQ) were very similar, sharing 92% identity (Figure 2A). For the Arf1/6 ancestral sequences, both programs predicted a large number of residues with high probability. 165/182 positions (90.66%) in the Arf1/6 IQ-TREE ancestor and 167/182 positions (91.76%) in the Arf1/6 PAML ancestor reconstructed with greater than 0.8 probability (Supplementary Table 2). Of those positions falling below this cut off, 10/17 in the Arf1/6IQ sequence and 7/15 in the Arf1/6PAML sequence conserved the basic chemical properties of the residue (Figure 2A). Notably, in seven cases, the PAML sequence included, as the residue reconstructed, the residue that was the second most likely from the IQ-TREE reconstruction. Reciprocally, there were four instances where the IQ-TREE sequence included the second most likely PAML residue at the same position. Thus we are confident that both the IQ-TREE and PAML reconstructions are robust and that together they cover the sequence space for possible ancestral reconstruction of the Arf1/6 ancestral protein (Figure 2A, Supplementary Table 2).

The ancestral Arf1/6 sequences predicted by PAML and IQ-TREE share a high level of identity to existing eukaryotic Arf1 and Arf6 sequences, especially in the nucleotide binding regions (G1 to G5, Figure 2B), which are highly conserved among Arf1 and Arf6 proteins themselves. The ancestral Arf1/6 sequences were more similar to Arf1 than to Arf6 proteins (Figure 2B). For example, Arf1/6PAML and Arf1/6IQ were 84% and 89% identical to human Arf1, and 73% and 72% identical to human Arf6, respectively. There was less conservation compared to individual eukaryotic Arf1 and Arf6 sequences in the N-terminal amphipathic helical (AH) region, which among eukaryotic sequences is also quite variable. However, the feature of this region that is important for Arf protein function is its amphipathic nature, which mediates tight membrane binding of the active GTP-bound form of the protein^34,35^. The AHs of the Arf1/6 predicted ancestral sequences clearly have this amphipathic property (Figure 2C). In addition, both Arf1 and Arf6 are myristoylated on the glycine residue following the initiating methionine, and both predicted ancestral sequences have this feature (Figure 2B, C).

The C-terminal regions of Arf1 and Arf6 proteins across eukaryotes are quite variable, both in length and amino acid composition, likely explaining the lower confidence levels of some C-terminal residues in the predicted ancestral sequences in this region (Figure 2B, Supplementary Table 2). Because of the length variation observed at the C-terminus of Arf1 and Arf6 sequences across eukaryotes, the positions corresponding to position 182 in the initial predictions could be included or excluded as a discretionary decision. We chose to test predicted sequences that were 181 and 182 amino acids in length, with the latter having an additional lysine residue at the C-terminus.

### Localization of Predicted Ancestral Proteins

In order to test the properties of the predicted ancestral proteins in cells, we chose two model eukaryotic systems, mammalian (MDCK; Madin-Darby Canine Kidney epithelial) cells and budding yeast (*Saccharomyces cerevisiae*). Starting with MDCK cells, we expressed Arf1/Arf6 ancestral proteins predicted by both the PAML and IQ-TREE programs, testing two predicted versions by each program, as described in the previous section. All four predicted proteins were fused to GFP at their C-termini, exogenously expressed using transient transfection, and compared to human Arf1 and Arf6 expressed from the same vector. Western blot, using anti-GFP antibodies, and in-gel GFP fluorescence analyses indicated similar levels of expression in mammalian cells (Supplementary Figure 3). The predicted ancestral proteins localized to the Golgi, as did Arf1, colocalizing with the Golgi marker GM130 (Figure 3). In contrast to Arf1, both Arf1/6PAML and Arf1/6IQ also localized to the PM, in a manner similar to Arf6, colocalizing with the 8+ PM probe (Figures 4 and 5). The major difference between the predicted ancestors and Arf6 that we observed is that the cytosolic pool of the Arf1/6 predicted ancestral proteins was less than that of Arf6, due to the fact that a large fraction of Arf1/6PAML and Arf1/6IQ also localized to the Golgi, unlike the case for Arf6. We found that the two versions of each predicted ancestral protein, differing in the number of C-terminal residues, had identical localization properties (Figure 5, Supplementary Figure 4). Thus, the predicted ancestral Arf1/6 proteins were found to specifically localize to the two membranes most commonly associated with Arf1 (Golgi) and Arf6 (PM).

**Figure 3:**
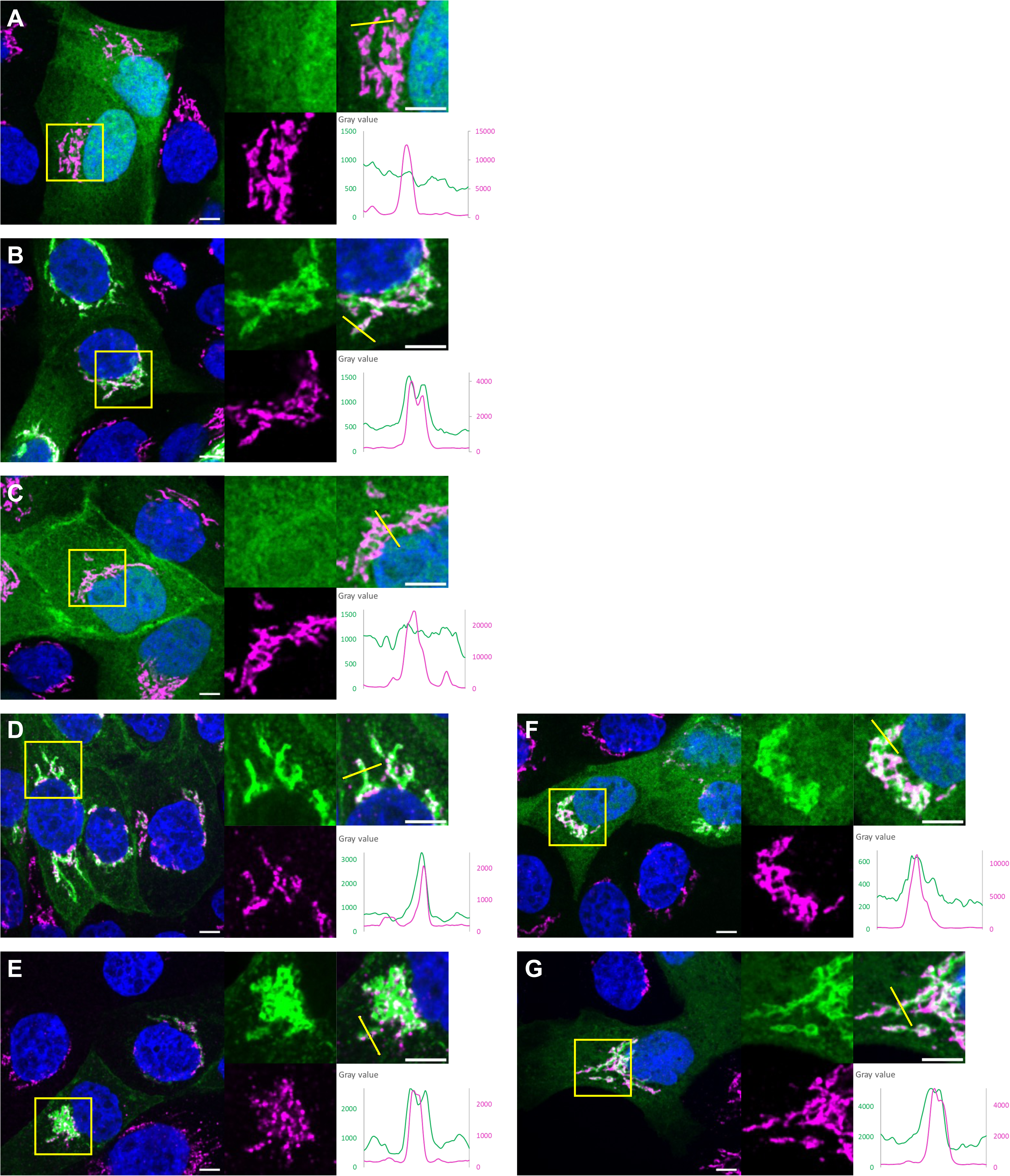
Localization of Arf1/6 predicted ancestral proteins to the Golgi in MDCK cells. MDCK cells were transfected with GFP (A); Arf1-GFP (B); Arf6-GFP (C); ARF1/6PAML-GFP (D); ARF1/6IQ-GFP (E); ARF1/6PAML-3KQ-GFP (F); or ARF1/6IQ-4KQ-GFP (G), and prepared for immunofluorescence analysis using antibodies against GM130, an early Golgi marker (magenta) and Hoechst dye to visualize nuclei (blue). Representative merged images are shown on the left for each transformant, with GFP- tagged protein (green), Golgi (magenta) and nuclei (blue). Insets (yellow square) are shown on the right, with the GFP and Golgi channels shown separately (left inset panels). The yellow lines shown in the merged inset panels (upper right) are those used to generate plots of pixel intensities for the GFP-tagged proteins (green) and the GM130 Golgi marker (magenta) (lower right) of each part (A-G). At least three independent transformations were performed for each mutant plasmid, always including the appropriate control plasmids. Scale bar, 5 μm.

**Figure 4:**
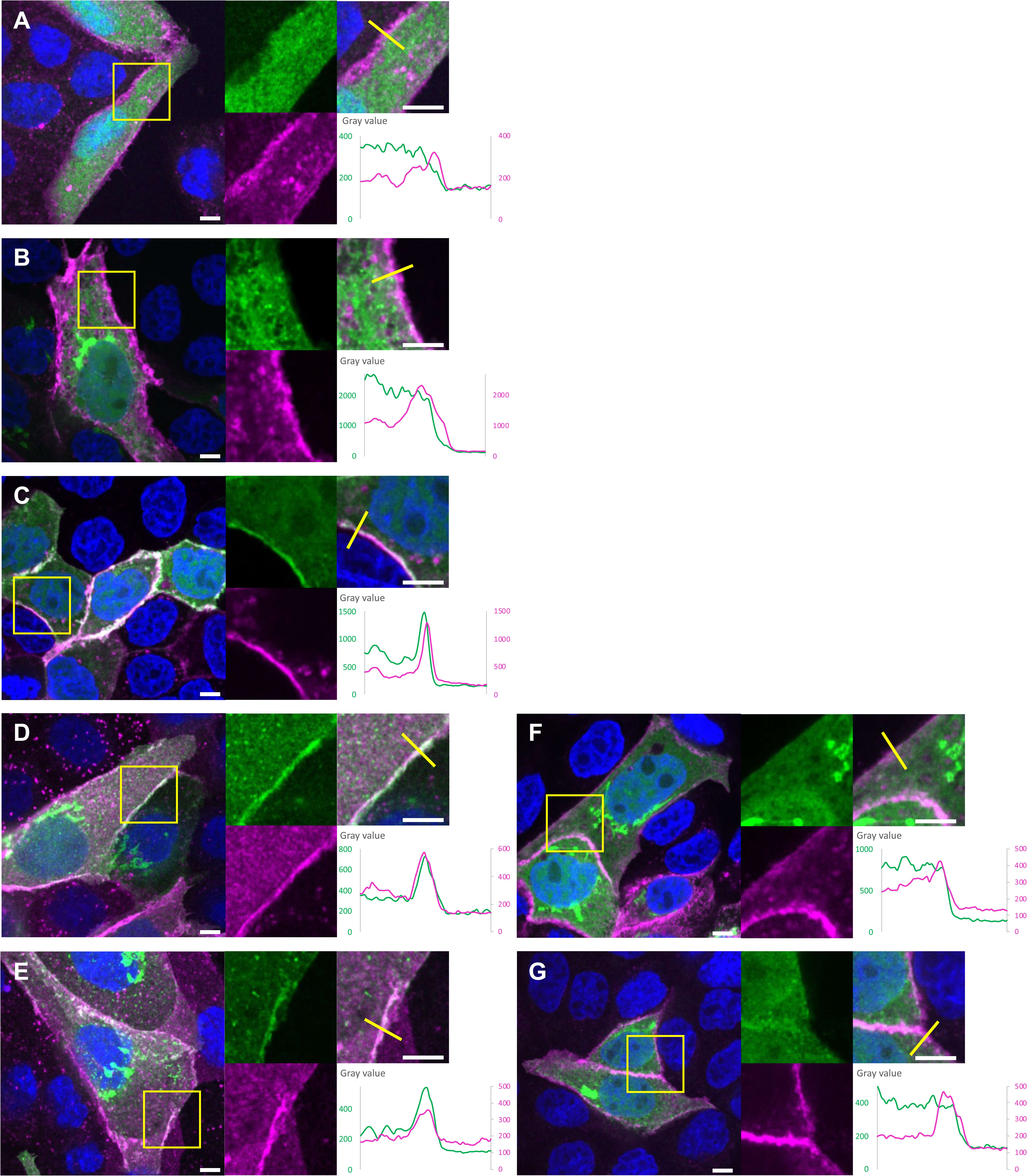
Localization of Arf1/6 predicted ancestral proteins to the PM in MDCK cells. MDCK cells were co-transfected with GFP (A); Arf1-GFP (B); Arf6-GFP (C); ARF1/6PAML-GFP (D); ARF1/6IQ-GFP (E); ARF1/6PAML-3KQ-GFP (F); or ARF1/6IQ-4KQ-GFP (G), and the 8+mCherry PM probe, then prepared for immunofluorescence analysis using Hoechst dye to visualize nuclei (blue). Representative merged images are shown on the left for each transformant, with GFP-tagged protein (green), PM (magenta) and nuclei (blue). Insets (yellow square) are shown on the right, with the GFP and PM channels shown separately (left inset panels). The yellow lines shown in the merged inset panels (upper right) are those used to generate plots of pixel intensities for the GFP-tagged proteins (green) and the PM marker (magenta) (lower right) of each part (A-G). At least three independent transformations were performed for each mutant plasmid, always including the appropriate control plasmids. Scale bar, 5 μm.

**Figure 5:**
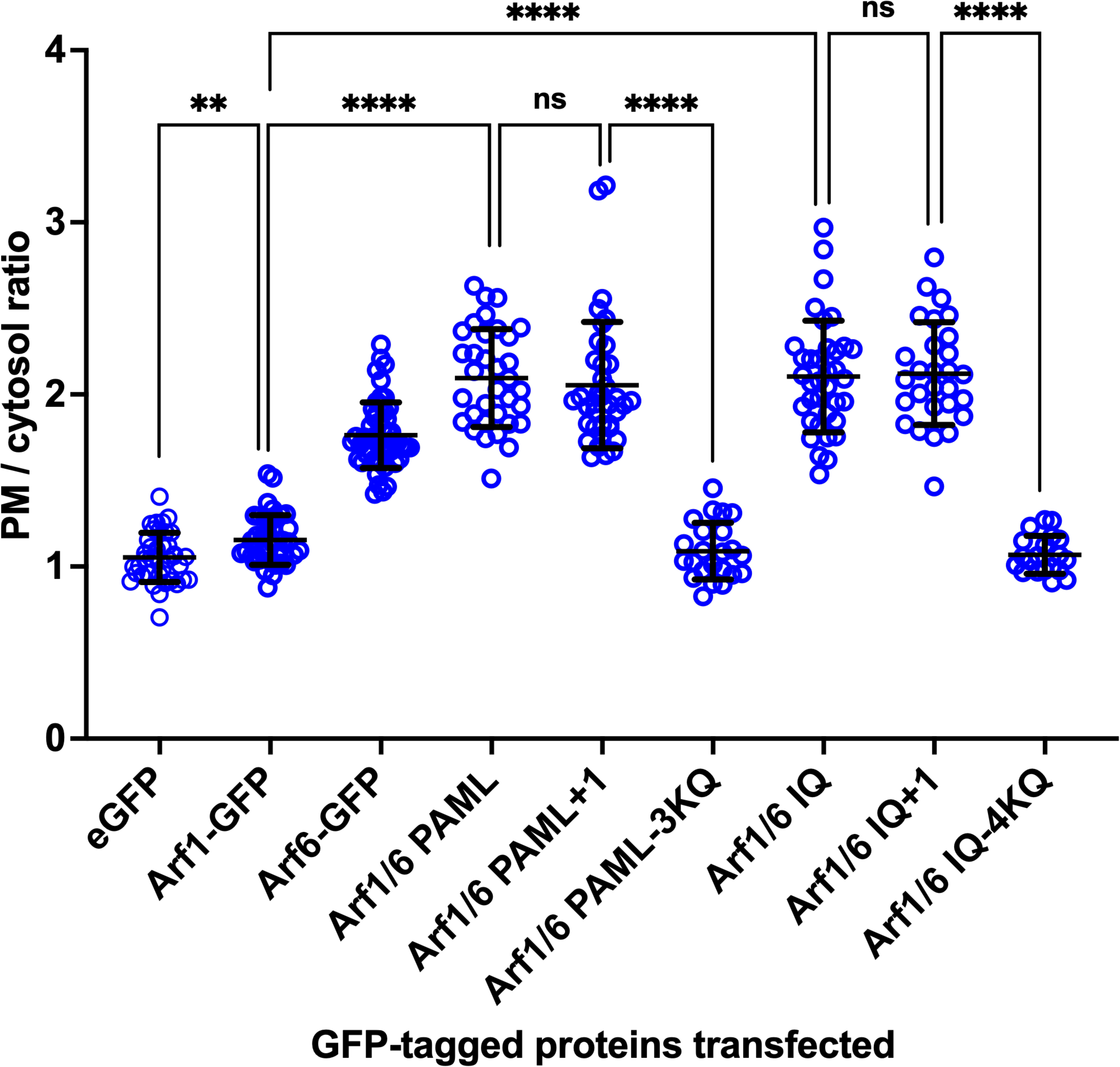
Quantification of PM localization. Levels of GFP-tagged proteins at the PM in individual cells (n) from the experiments shown in Figure 4 and Supplementary Figure 4 were quantified. For GFP, n=30; Arf1-GFP, n=30; Arf6-GFP, n=15; ARF1/6PAML-GFP, n=33; ARF1/6IQ-GFP, n=19; ARF1/6PAML+1-GFP, n=18; ARF1/6IQ+1-GFP, n=22; ARF1/6PAML-3KQ-GFP, n=18; ARF1/6IQ-4KQ-GFP, n=18. The membrane/cytosol ratio was calculated by dividing the membrane fluorescence value by the average cytosolic fluorescence value, as described in Methods. Data are presented as mean values +/- standard deviation, with each individual point shown. Statistical analysis (unpaired t test with Welch’s correction, two-tailed p value) was performed. ** indicates p < 0.01; ****, p < 0.0001; ns, not significant. p = for the comparison of GFP alone and Arf1-GFP. Numerical values obtained for all calculated ratios are in Supplementary Table 4, and p values for all pairwise comparisons are in Supplementary Table 5.

We also tested localization of the Arf1/6PAML and Arf1/6IQ ancestral proteins in yeast cells. Western blot analysis indicated similar levels of expression of the recombinant, GFP-tagged proteins in yeast (data not shown). The yeast Golgi is not a single, centrosomally localized organelle as in mammalian cells, but is fragmented into individual elements dispersed throughout the cytosol. We used yeast Arf1 fused to mCherry as a Golgi marker, as the majority of Arf1 at steady state is present at the Golgi, colocalizing with the early Golgi marker α-COP (Cop1), part of the COPI coat, and with the late Golgi marker clathrin heavy chain (Chc1) ^36^. We found a high level of colocalization of both Arf1/6 predicted ancestral proteins with yeast Arf1 (Figure 6). In addition, unlike yeast Arf1, both Arf1/6PAML and Arf1/6IQ were also found at the PM (Figure 7). As was the case in mammalian cells, we found that the two versions of each predicted ancestral protein, differing in the number of C-terminal residues, had identical localization properties (Supplementary Figure 5). Hence in two different eukaryotic cellular systems, with quite distinct organellar morphologies, the Arf1/6 predicted ancestral proteins have localization properties of both Arf1 and Arf6, localizing to both the Golgi and the PM, suggesting that the common ancestor was able to localize to both of these compartments.

**Figure 6:**
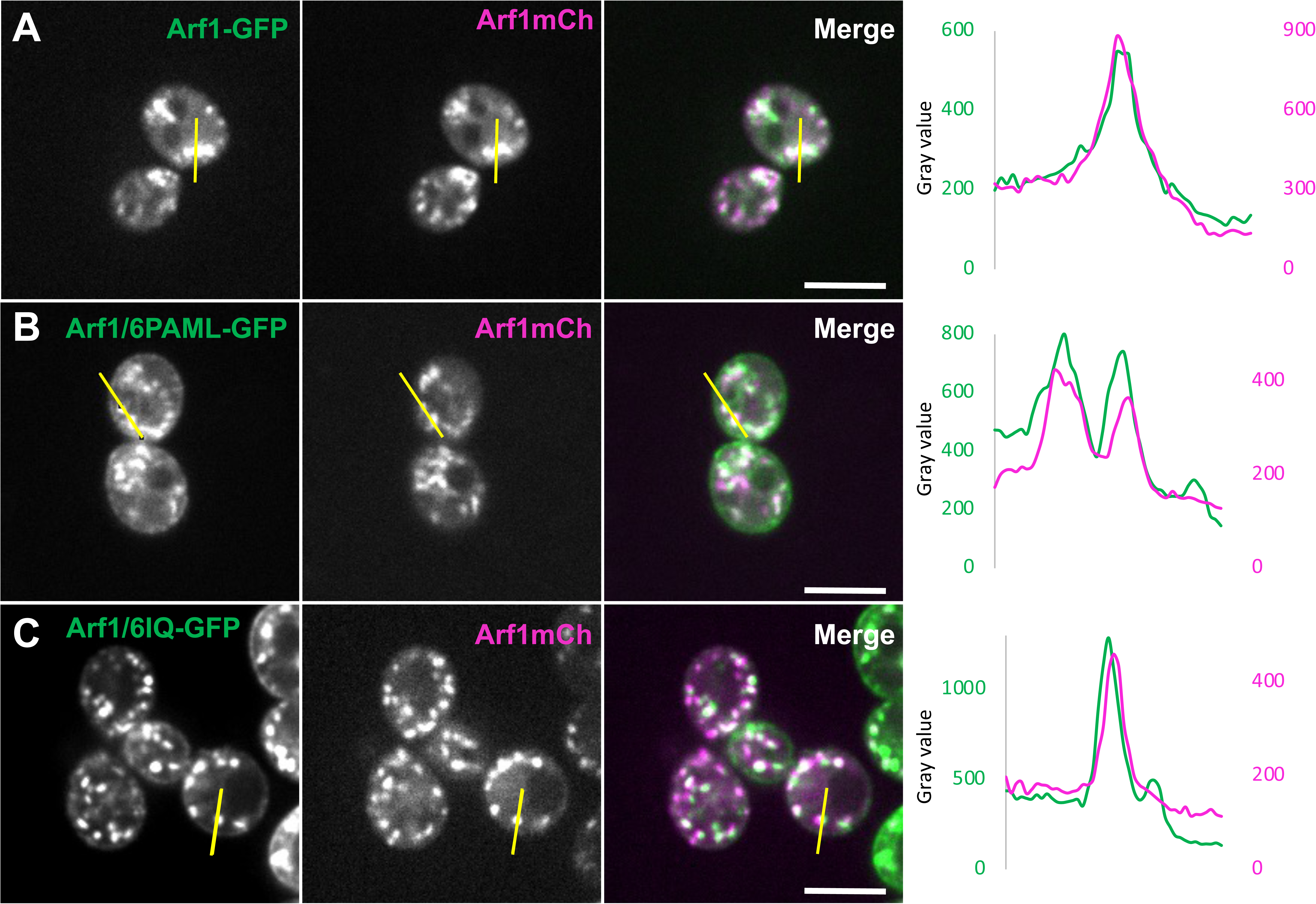
Localization of Arf1/6 predicted ancestral proteins in yeast cells. Plasmids encoding Arf1-GFP (A), Arf1/6PAML-GFP (B) and Arf1/6IQ-GFP (C) were co-transformed into *Saccharomyces cerevisiae* cells with a plasmid encoding Arf1-mCherry. Representative images for each GFP-tagged protein expressed (left panels), with Arf1-mCherry (middle panels) are shown. The yellow lines are those used to generate plots of pixel intensities (right panels) for the GFP-tagged proteins (green) and Arf1-mCherry (magenta). At least three independent transformations were performed for each mutant plasmid, always including the WT control plasmid. Scale bar, 5 μm.

**Figure 7:**
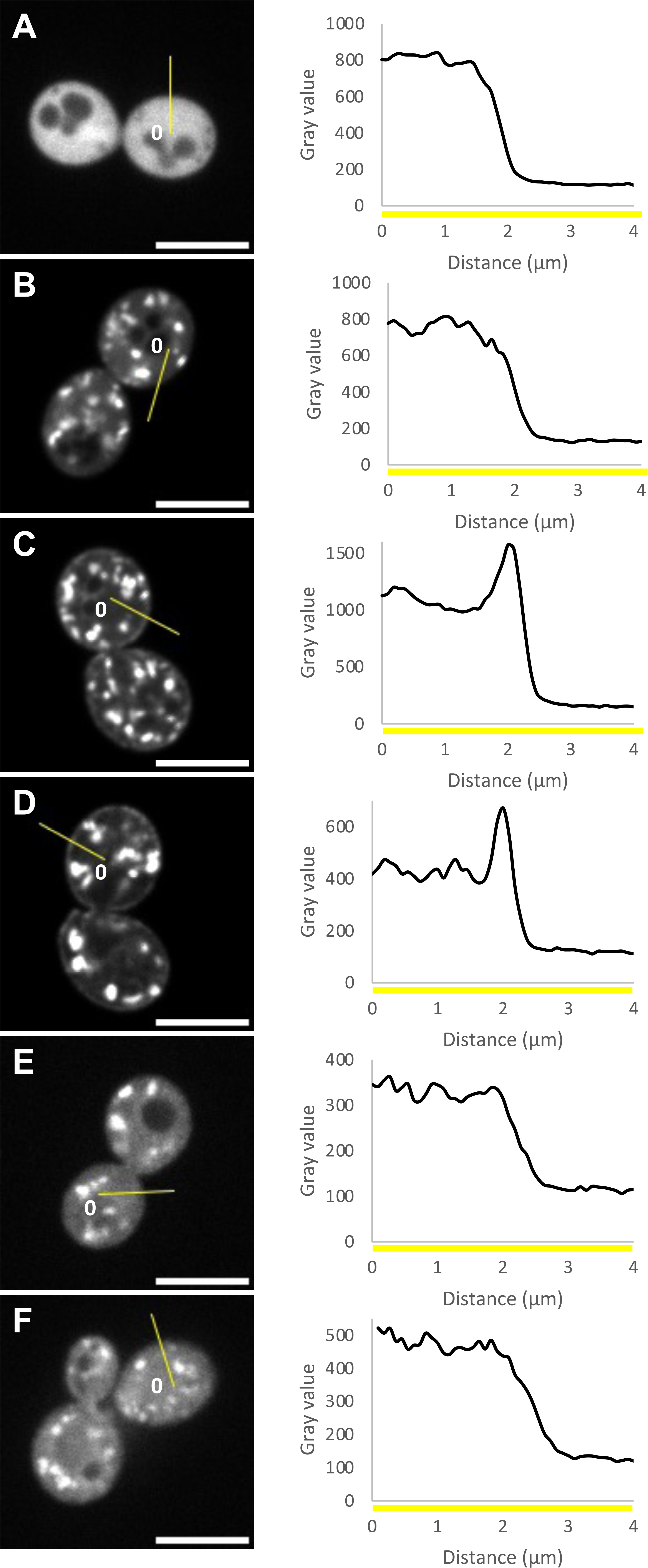
Localization of Arf1/6 predicted ancestral proteins to the PM in yeast cells. Plasmids encoding GFP (A), Arf1-GFP (B), Arf1/6PAML-GFP (C), Arf1/6IQ-GFP (D), Arf1/6PAML-3KQ-GFP (E), and Arf1/6IQ-4KQ-GFP (F) were transformed into *Saccharomyces cerevisiae* cells. Localizations of each GFP-tagged protein were quantified as described in Methods. Representative images for each protein expressed in yeast are shown (left panels). The yellow lines are those used to generate plots of pixel intensities (right panels). “0” (left panels) indicates the beginning of the line shown, and corresponds to “0” on the x-axis of the plots (left panels). At least three independent transformations were performed for each mutant plasmid, always including the WT control plasmid. Scale bar, 5 μm.

### Mechanism of Localization

Previous work has shown that one localization determinant for the Golgi in Arf1 is present in the alpha3 helix and downstream residues (aa 101 to 116 in *Homo sapiens* and *Arabidopsis thaliana* Arf1), and contains the MXXE motif^23,24,37^. This motif is lost in Arf6 proteins. However, when these amino acids are re-introduced into human Arf6, they are sufficient to target Arf6 to the Golgi^23^. We found that the predicted ancestral proteins have the MXXE motif, consistent with their Golgi localization. We also noted that the predicted ancestral proteins have multiple lysines at their C-termini, a known PM localization determinant that binds specifically to anionic lipids enriched in the PM^38–40^. We mutated these lysine residues to glutamines, and expressed the mutants (Arf1/6PAML-3KQ and Arf1/6IQ-4KQ) in mammalian cells. For both Arf1/6PAML and Arf1/6IQ predicted ancestral sequences, the mutant versions failed to localize to the PM (Figures 4 and 5). However, we observed Golgi localization for these C-terminal KQ mutants. Similarly, when the C-terminal KQ mutants of both Arf1/6PAML and Arf1/6IQ sequences were expressed in yeast cells, localization to the PM was lost, but localization to the Golgi was unaffected (Figure 7). We conclude that a property of the Arf1/6 predicted ancestral protein is that it had the capacity to localize to both the Golgi and the PM, and that PM localization is mediated by C-terminal basic residues.

### Diverse Modern Arf1 Homologues Possess C-terminal Lysines

Because Arf6 localizes and functions at the PM, we expected that the PM localization of the Arf1/6 ancestor has its basis in Arf6 sequences. Indeed, the Arf6 predicted ancestral sequences do have multiple C-terminal basic residues (Supplementary Table 2). However, surprisingly, the predicted Arf1 ancestral sequences (using either PAML or IQ-TREE) also have multiple C-terminal positively charged residues (Supplementary Table 2). Examination of the Arf1 sequences used in the ASR indicates that 31 of 120 Arf1 sequences in extant eukaryotes have multiple C-terminal, basic residues. Moreover, the Arl1 (11 of 108) and Arl5 orthologues (25 of 79) also possess the C-terminal basic residue motif. By contrast, only 1 of 55 orthologues of Arf6 included in the ASR dataset possesses this motif. Thus, and somewhat counter-intuitively, the inferred PM-localizing motif in the Arf1/6 reconstructed ancestral proteins does not arise from a highly prevalent motif found in Arf6 orthlogues used in this analysis, Rather it is due to the more frequent presence of the motif in the other three subfamilies in our dataset, and thus calculated to be an ancestral feature of each.

To further explore the prevalence of the polybasic C-terminal region in modern Arf1 proteins, we examined the broad diversity of eukaryotic Arf1 sequences in our more extensively sampled dataset, a subset of which was used for the ASR. From a highly conserved W (aa 172 in human Arf1) onwards, we examined sequences to determine the number of basic residues. We found that Arf1 orthologues with positively charged residues in the C-terminus are found all across the eukaryotic tree of life. In humans, the Arf1 orthologue Arf3 has 3 C-terminal lysines, and has been shown to localize to more peripheral structures thant Arf1 and to function in plasma membrane processes^41^. A selection of Arf1 sequences with multiple basic residues, from a range of eukaryotes, and human Arf1 for comparison, is shown in Figure 8, and includes organisms of medical, and ecological impact. Notably, in addition to those Arf1 sequences with C-terminal basic residues, there were many cases in which a given taxon possessed multiple Arf1 homologues, and that used for the ASR was not the one with the highest number of C-terminal basic residues. This means that if anything the ASR analysis represents an under-estimate of the basic residue count in the Arf1/6 ancestral sequences. It also means that the C-terminal basic tail, and by inference the PM localization of Arf1, is a more prevalent feature of modern eukaryotic Arf1 proteins than currently understood.

**Figure 8:**
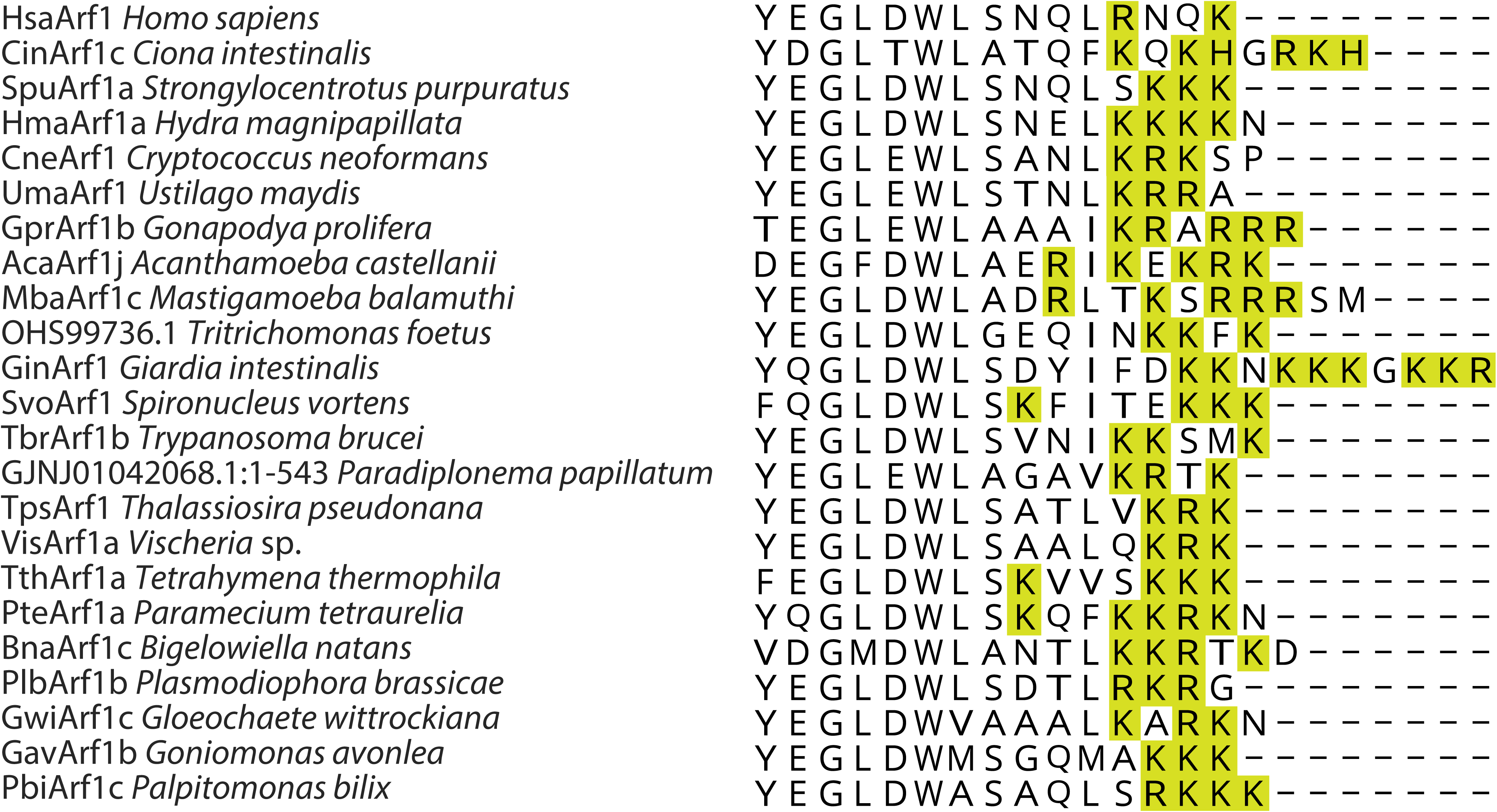
C-terminal region of selected Arf1 homologues from diverse eukaryotes. This alignment illustrates the presence of multiple positively charged residues in the C-terminus of Arf1 proteins from phylogenetically diverse eukaryotes, including animals, fungi, and organisms with agricultural (*P. brassicae*, *T. foetus*), ecological (*T. pseudonana*, *P. papillatum*), and medical (*G. intestinalis*, *C. neoformans*, *T. brucei*) impact. The sequence abbreviation used in Supplementary Table 1 is given before the organism name. Positively charged residues are coloured lime.

## Discussion

In this work we explored the cell biological properties of the ancestor of Arf1 and Arf6, a protein which predated LECA, by characterizing four predicted ancestral sequences. In the process of addressing this question of ancient evolution, we learned new properties of existing Arf1 and Arf6 proteins in a wide range of eukaryotes. Arf family proteins are key regulators of membrane traffic, and are ancient, having arisen in the archaeal ancestor of eukaryotes ^8,25^. These results place the Arf family GTPases at the very beginning of the process of eukaryogenesis. Between the archaeal ancestor and LECA, expansion of the Arf family proteins occurred, leading to the LECA complement of 15 members of this family, including Arf1 and Arf6 ^12^. In well-studied eukaryotic systems, the major localization of Arf1 is at the Golgi, where a highly conserved function is in vesicular traffic from the Golgi to either the ER or the endosomal system ^27,42,43^, whereas Arf6 localizes predominantly to the PM and endosomes, where it regulates the actin cytoskeleton and endocytic pathways ^13,14,26^. Although having primary functions at different organelles, Arf1 and Arf6 are similar at the sequence level, even sharing many amino acids in the switch regions, which interact with effectors in the GTP-bound state^44–49^. Indeed, localization of mammalian Arf1 and Arf6 to different membrane sites is thought to be a major factor in their distinct functions. For all of these reasons, the Arf1/6 ancestor is a theoretical reconstruction point particularly well suited to the APR approach of addressing pre-LECA evolution.

We found that the predicted Arf1/6 ancestors are far more similar to Arf1 than to Arf6 proteins at the primary sequence level, and that these ancestors had the capacity to target the PM using a C-terminal polybasic motif. Unexpectedly, this motif is also found in a number of existing Arf1 proteins (and also in GTPases of the Arl1 and Arl5 outgroups), presumably retained from the ancestral form. Many of the Arf1 proteins we identified with C-terminal polybasic motifs are present in organisms for which little or no localization or functional data are available. However, studies have been carried out on the Trypanosoma brucei Arf1 protein TbArf1, which has clearly been shown to be an orthologue of Arf1^12^, and which has three C-terminal K residues (Figure 8). TbArf1 localizes to the Golgi but also has endocytic functions in Trypanosoma cells that are normally regulated by Arf6 in mammalian and yeast cells. Notably, in addition, TbArf1 has a strong PM localization when expressed in mammalian cells^50^.

It is worth considering the limitations of our work and clarifying the steps that were taken to mitigate these factors. While ASR is of course a theoretical reconstruction, we took several steps to provide the most reasonable reconstruction and avoid biases. We sampled organisms from across the broad diversity of eukaryotes, providing a balanced sampling. We performed upstream analyses to select the least divergent representatives and we used two different ASR methods, each with sophisticated models of sequence evolution. The results of these analyses showed overall conservation with well-characterized Arf proteins (Figure 2). In fact, these were nearly identical in some cases to Arf proteins found in existing organisms (e.g. *Planomonas micra*), which lie near the most recent predicted position of the root of the eukaryotic tree ^51^. From an evolutionary perspective, Arf1 and Arf6 are one another’s closest relatives in an unrooted tree of Arf family proteins^12^, as compared to other possible reconstructions where there are other protein subfamilies intervening. We did assume that the root of the Arf tree does not lie on Arf1, Arf6 or basally to Arf1/6, thus making the Arl1/5 clade an outgroup. Although this is a declared assumption, we believe that it is a strong one. Past attempts to root the Arf family tree with the recently described ArfR archael homologues were unresolved but gave no indication of a root on the Arf1, Arf6, or Arf1/6 groups ^8^. Furthermore, a rooting on Arf1, Arf6 or Arf1/6 would then imply that all other Arf family sub-families are derived from Arfs and evolved at a later evolutionary point. This would include the COPII GTPase Sar1. We believe this is unlikely, particularly as other components of the COPII machinery (e.g. Sec23/24) homologues have been identified in Asgardarchaea ^7^. Finally, recognizing the theoretical nature of our reconstructions, we have limited our conclusions to broad properties of the sequences, rather than specific amino acids, and corroborated with experimental data or sequences from existing organisms. Even with the steps taken for robustness in our predictions, we chose to test sequences derived from both methods to cover the range of sequences that we had reconstructed and balanced against methodological bias. We also chose to perform our analyses in two model organisms, those that are the best studied and from which a large proportion of the biological knowledge about Arf family proteins have been determined^13,14,27,52,53^. Importantly, we observed experimental consistency in the two cell biological model systems and in the predicted ancestral sequences reconstructed using both methods. With all of these efforts for robustness, we believe that there are important conclusions that can be drawn.

Arf1 and Arf6 are localized to membranes through multiple mechanisms, including recruitment of the GDP-bound form through a membrane-bound receptor, interaction with specifically localized Arf guanine nucleotide exchange factors (GEFs) and by activation and interaction with effectors. The complete mechanism of localization for any one of these GTPases has not been fully resolved ^27^. Although in vitro, Arf1 and Arf6 are soluble in their GDP-bound forms, in cells the GDP-bound forms of both GTPases have been shown to bind to membranes via protein-protein interactions. Arf6 in its GDP-bound form localizes to the PM^18^ and interacts specifically with the TBC domain proteins TRE17 and TBC1D24/TBC-K, mediating PM association^17,19^. Arf1 has been shown to bind to at least three Golgi proteins in its GDP-bound form, the Golgi-localized SNARE protein membrin^23,24^ and members of the p24 family^20,21^ in mammalian cells, and to members of the ArfGAP2/ArfGAP3/Glo3 family of ArfGAPs in plant cells^22^.

The interaction of Arf1-GDP with membrin is the best studied, and has been shown to mediate specific localization of the Arf1 GTPase to the Golgi^23,24,37^. The region of Arf1-GDP that interacts with membrin, the alpha3 helix and four residues downstream, is sufficient for Golgi localization, and has a markedly different sequence in Arf6^23^. The importance for Golgi localization of the MXXE motif within this region was first demonstrated in mammalian cells^23^ and was also shown for a plant Arf1 orthologue^24,37^. Indeed, mutagenesis studies showed that mutations in the M110 and E113 residues within the MXXE motif of both *Homo sapiens* and *Arabidopsis thaliana* Arf1 reduce Golgi localization^23,37^. Our results show that this feature is widespread throughout eukaryotic Arf1 orthologues, but is not found in Arf6 sequences. Our studies predict that it was present in the ancestor of Arf1 and Arf6 proteins.

Even though Arf6 proteins in many organisms including mammals and yeast do not have a C-terminal polybasic motif, there is one known mechanism of Arf6 localization to the PM that does depend on one. This mechanism is via an Arf cascade in metazoan cells, whereby PM-localized Arl4 recruits the Arf6 activator (GEF) cytohesin, which in turn activates and recruits Arf6 ^54–56^. There are three Arl4 proteins in mammalian cells (Arl4A, C, D), all of which have nine lysine and/or arginine residues at their C-termini that are required for PM localization through binding to anionic PM lipids such as PI(4,5)P_2_ ^40,54^. This Arl4-dependent mechanism of Arf6 localization was established late in evolution, as Arl4 proteins arose in early metazoans, likely from Arl19. Arl19 arose in the stem branch of Holozoa, possibly from Arf1 ^16^, and numerous Arl19 orthologues also have multiple basic residues at their C-termini ^12^.

This work has important implications for understanding both the cell biology of Arf proteins in modern eukaryotes and the process of eukaryogenesis. Our results predict that the eukaryotic ancestor of Arf1 and Arf6 proteins localized to the Golgi and the PM via mechanisms that were maintained in many modern lineages. Arf1 paralogues with a poly-basic C-terminal motif are present in diverse eukaryotes including mammalian cells and important parasites of global health importance (e.g. *Giardia intestinalis*), economically relevant agricultural or veterinary pathogens (e.g. *Tritrichomonas foetus*, *Ustilago maydis*, *Plasmodiophora brassicae*) or protists with key ecological roles (e.g. the diatom *Thallasiosira pseudonana,* the dinoflagellate *Symbiodinium minutum,* or the heterotrophic flagellate *Paradiplonema papillatum*). Hence for a broad range of eukaryotes, we predict that some of their Arf1 proteins will not follow the current paradigm for Arf1 and will localize to the cell periphery, as well as the Golgi. Indeed, in mammalian cells, the idea that Arf1 proteins have broader functions than at the Golgi has been gaining support, where Arf1 has been shown to have functions at the PM, for example in phagocytosis, cell migration, neutrophil chemotaxis, and the CLIC/GEEC endocytic pathway ^57–63^. In addition, Arf6 itself can recruit cytohesin which then recruits Arf1 to the PM in another Arf activation cascade ^46,47^. This mechanism enables a strong and sustained activation signal at the PM, as Arf1 is more abundant than either Arf6 or Arl4 ^47^. Our results support the conclusion that the multifunctionality of Arf1 is an evolutionarily conserved feature.

With respect to the events and processes of eukaryogenesis, our results strongly suggest that the Arf1/6 ancestor was a generalist protein acting at multiple cellular membranes in pre-LECA organisms. Based on the observed distribution of subfamily orthologues with the C-terminal basic motif, and the presumed rooting outside the Arf1/6+Arf1/6 clades, the most straightforward evolutionary scenario is that the ancestor of Arf1, Arf6, Arl1 and Arl5 also likely acted at multiple locations. This feature was retained in the ancestor of each subfamily. This mechanism was either retained, lost (and thus subfunctionalized to a specific cellular location) or replaced, as in the Arl19 mechanism, in the respective paralogues of the subfamily members in descendant eukaryotic lineages after LECA. Future testing of this scenario will hinge on a resolved rooting for the entire Arf family tree and on experimental characterization of Arf1, Arf6, Arl1 and Arl5 proteins in diverse eukaryotes.

While ancient protein resurrection has been used to investigate deep nodes within the eukaryotic tree ^31^ and even aspects of LECA proteins ^30^, to our knowledge, this is the first attempt to reconstruct a pre-LECA protein. In this case it appears that a multifunctional protein duplicated, with one paralogue retaining its generalist nature and the other paralogue undergoing divergence and loss in many cases. Loss after duplication is a common result but long-term retention, divergence, and sporadic loss over a 1.5 billion year span suggests complex cell biological involvement. This requires a much deeper understanding of Arf6 function to explain, particularly in non-opisthokont organisms where cell biological information is lacking. Nonetheless, our work does demonstrate that the ASR approach to understanding events in the intervening eukaryogenic period between first eukaryotic common ancestor (FECA) and LECA is tractable and can yield concrete results. We contend that it opens up a highly promising new avenue of investigation for an important and currently intransigent problem in evolutionary biology.

## Methods

### Homology Searching

Our sampling was build upon that from Vargová et al. 2021 ^12^. Nonetheless, to incorporate sampling from newly sequenced or described taxonomic lineages (e.g. ancyromonads, CRuMs, Telonemia), we searched 24 additional ‘omics dataset (Supplementary Table 1). Human Arf1, Arf6, Arl1, and Arl5 were used as the query sequences to search into the dataset for Arf family homologs with the human genome being used as the reference genome.

Homology searching for the Arf family paralogs were carried out with Analysis of MOlecular Evolution with BAtch Entry (AMOEBAE) ^64^. Forward searches into the assembled database were carried out with BLASTp to identify potential Arf homologs that have sufficient E-value support(E<0.05). Potential homologues were used as queries to search into the reference genome to check if confirmed Arf family subunits are also detected with proper E value support (E<0.05 and must be a factor of 10^2^ less than next nonredundant hit). Sequences that were clearly fragmentary were discarded. The remaining data was combined with a selection from Vargová et al. 2021 ^12^.

### Phylogenetic Analyses

For computational tractability, the dataset was then subdivided into subdatasets for Arf1, Arf6, Arl1, and Arl5. Alignment was performed with MUSCLE (V5.1) and masked/trimmed manually using Mesquite (V3.70). IQ-TREE (V2.2) was run for 1,000 ultrafast bootstraps for each of the Arf subfamilies with default model finding criteria using the embedded Model finder (analysis details in legends of Supplementary Figure 1A-D). The least divergent paralogue (shortest branch length) for each of the Arf families analyzed was collected for further analysis for a total of 120 Arf1, 56 Arf6, 108 Arl1, and 79 Arl5 homologues across a total of 120 species and an alignment of 182 position.

C-terminal basic residue motif assessment was based on a visual inspection of the above alignments. If a sequence possessed three or more basic residues (Arginine, Lysine, Histidine) present after the highly conserved Tryptophan (aa 172 in human Arf1) onwards, it was deemed to be affirmative for the presence of the motif. In some cases sequences were omitted from the total count if it was clear that they had been truncated at the C-terminus prior to W172.

### Ancestral Sequence Reconstruction

The datasets were recombined and aligned using MAFFT and manually trimmed to remove unique indels. An unresolved guide tree was imposed, following the most up to date topology of the eukaryotic tree of life ^32,33^ and respecting the sister clade relationship of Arf1/Arf6 and Arl1/Arl5 ^12^. The Arf1 and Arf6 tree was calculated with a rooted constraint of the Arl1/Arf5 clades that are the closest sister clades in an unrooted phylogeny from Vargová et al. 2021^12^ as an assumed outgroup.

Ancestral sequence reconstruction using the constrained guidetree and an alignment of eukaryotic Arfs was performed using an empirical (model=2) non mixture LG model on Phylogenetic Analysis by Maximum Likelihood (PAML V4.8) based on the results of IQ-TREE ModelFinder. Ancestral sequence reconstruction was also performed using a LG+C60+R8 mixture model in IQ-TREE, derived from a ModelFinder search for the most appropriate mixture model. The ASR was also calculated for the Arf1/6 ancestors. Because of the length variation observed at the C-terminus across eukaryotes, the positions corresponding to positions 182-183 could be included or excluded as a discretionary decision. To be conservative, we eliminated the positions, resulting in final sequences of the Arf1 ancestor (181 positions in both IQ-TREE and PAML) ending with a chain of three lysine residues and the Arf1/6 ancestor (181 positions in both IQ-TREE and PAML) with four lysines (Figure 2).

The N-terminal AHs of the PAML and IQ-TREE predicted ancestral sequences were determined from alignment with human Arf1 and Arf6, whose AH regions have been determined by crystal structure analysis ^65,66^. Helical wheel plots were generated by Heliquest ^67^.

### Cell Culture and Plasmids

MDCK cells (ATCC CCL-34) were grown in Dulbecco’s modified Eagle’s medium (DMEM) (Gibco #31885023) supplemented with 10% FBS (Gibco #A5256701) and 1% penicillin-streptomycin (Gibco #15140122). For transfections, cells were plated at 50% confluency in 6- or 24-well plates, and containing 1.5 mm coverslips for immunofluorescence microscopy. After 24 hours cells were transfected with 1 µL/cm^2^ of Lipofectamine 2000 (Thermofisher #11668027) and a total of 0.5 µg/cm^2^ of DNA, including plasmids of interest and carrier DNA where appropriate. After 48 hours, cells were fixed and prepared for immunofluorescence microscopy, or cell lysates were prepared either for western blotting or for in gel fluorescence (IGF) using a ChemiDoc MP imager (Bio-Rad) ^68^. As a loading control, total proteins were visualized in commercial pre-cast SDS–PAGE gels (stain-free TGX gels, 4%–20%; Bio-Rad) after photoactivation induced by a 45 second UV treatment using a ChemiDoc MP imager (Bio-Rad).

Yeast transformations were carried out using the lithium acetate method. Briefly, the BY4742 *MATα his3Δ1 leu2Δ0 lys2Δ0 ura3Δ0* strain^69^ was grown in YPD medium at 30°C to exponential phase, and approximately 4 OD_600_ equivalents of cells per transformation reaction were harvested, washed and resuspended in 60 µL of transformation solution 1 (10 mM Tris pH7.5, 100 mM LiAc, 1 mM EDTA). 2 µg of plasmid DNA and carrier DNA from salmon testes (Sigma #D9156-1ml, incubated at 95°C for 5 min), were added, then 300 µL of 40% PEG in transformation solution 1. Cells were incubated 30 min at 22°C or 30°C, then at 42°C for 15 min. Cells were resuspend in YPD, incubated 1 hour, then plated on selective medium. For live cell imaging, cells were grown in selective liquid medium, to a maximum OD_600_ of 0.3, concentrated, then observed by spinning disk microscopy. For western blotting analysis, cells were grown to a maximum OD_600_ of 0.5, and cell lysates prepared. After transfer of gels, nitrocellulose membranes were incubated with anti-GFP primary antibodies (Roche #11814460001) and anti-mouse IgG-HRP secondary antibodies (GE Healthcare, #NA931V). As a loading control, total proteins were visualized in commercial pre-cast SDS–PAGE stain-free gels after photoactivation induced by a 45 second UV treatment using a ChemiDoc MP imager (Bio-Rad).

Plasmids used in this study are listed in Supplementary Table 3. For mammalian cell exogenous expression, coding regions of Arf1, Arf6 and predicted ancestral proteins were cloned into the pEGFP-N3 vector, with proteins expressed under control of the human cytomegalovirus (CMV) immediate early promoter. The 8+mCherry PM probe was derived from the 8+GFP PM probe described previously ^38^. Yeast expression plasmids are centromeric vectors, with proteins under control of the constitutive ADH promoter.

### Immunofluorescence

The transfected MDCK cells previously seeded on 1.5 mm coverslips were fixed with 4% paraformaldehyde (PFA) for 15 min. PFA was inactivated with 50 mM NH_4_Cl for 20 min. After 2 1× PBS washes, cells were permeabilized for 30 min with 1× PBS, 0.025% saponin (Sigma-Merck #84510), then blocked with 1× PBS, 0.025% saponin, 1% BSA for 30 min. The primary mouse antibody targeting GM130 (BD Biosciences #610822) was hybridized in 1× PBS, 0.025% saponin, 1% BSA at 1:500 overnight at 4°C. After multiple washes, goat anti-mouse IgG Alexa fluor 647 (ThermoFisher #A-21235) was added at 1:500 for 1 hour. After multiple washes, nuclei were stained using Hoechst33342 (ThermoFisher #H3570) in 1× PBS at 1:10,000 for 15 min. Coverslips were washed again, rinced in water, then mounted using Fluoromount (ThermoFisher #00-4958-02). After 24 hours at room temperature, the cells were observed by confocal microscopy.

### Microscopy

For MDCK cells, acquisitions were performed either on a Zeiss confocal 980 Airyscan microscope driven by ZEN software (version 3.9.101.03000), using a 63× objective, or on a Leica Spinning Disk CSU-X1 microscope driven by MetaMorph software (version 7.10.4.407), using a 100× objective. For the images obtained using the Zeiss confocal 980 microscope, 40 z-sections separated by 0.13 µm were acquired for each field of cells using excitation/emission wavelengths 353/465 (Hoechst), 488/509 (GFP), and 587/610 (mCherry), and/or 653/668 (Alexa 647). For the Leica Spinning Disk microscope, 20 z-sections separated by 0.2 µm were acquired for each field of cells using excitation wavelengths of 405 (Hoechst), 488 (GFP), 561 (mCherry), 642 (Alexa 647), and emission filters 450/60 (Hoechst), 525/50 (GFP), 590/35 (mCherry), 670/40 (Alexa 647). Localization of GFP-tagged proteins and compartment markers was analysed in ImageJ2 V2.14.0/1.54f. Median and gaussian blur filters were applied to image stacks, and a projection of four z-sections was used for quantifications. A six-pixel wide line was drawn across a portion of the compartment of interest (PM or Golgi), identified by specific markers (8+mCherry or GM130, respectively). Examples are shown in Figures 3 and 4, yellow lines. Fluorescence intensity in each channel, expressed as a gray value, was obtained for each pixel along the line. Plots of fluorescence intensities across the yellow lines in the upper right panels of each part of Figures 3 and 4 are shown in the bottom right panels. To determine the membrane/cytosol ratio (Figure 5), the value of the peak of fluorescence at the PM was determined, and the average cytoplasolic fluorescence intensity was obtained by selection of a portion of the line passing over cytosol free of organelles, and averaging the fluorescence values obtained. The membrane/cytosol ratio for PM staining was calculated by dividing the membrane fluorescence value by the average cytosolic fluorescence value. For this quantification of PM localization, 20-53 cells from separate fields were analysed (numbers for each condition are given in Supplementary Table 4). The number “n” refers to the number of individual cells quantified for each plasmid transfected, chosen from different microscopy fields and from at least three different transfection experiments. Mean and standard deviation of the calculated membrane/cytosol ratios were obtained, and statistical analysis (unpaired t test with Welch’s correction, two-tailed p value) was performed, using Graphpad Prism software (V10.5.0). **** indicates p < 0.0001; **, p < 0.01; ns, not significant. The values of p for each pairwise comparison are given in Supplementary Table 5. Numerical values of the raw data obtained, as well as the results of statistical analyses are in Supplementary Table 4.

For yeast cells, acquisitions were performed on a Leica Spinning Disk CSU-X1 microscope, driven by Metamorph software, using a 100× objective. 21 z-sections separated by 0.2 µm were acquired for each field of cells, using excitation wavelengths of 488 (GFP) and 561 (mCherry), and emission filters 525/50 (GFP) and 590/35 (mCherry). Localization of GFP-tagged proteins was analysed using ImageJ2 by selecting an appropriate z-section, choosing cells with well-defined organelle localization, and drawing a line across the cell (examples are shown in Figures 6 and 7, yellow lines). Fluorescence intensity was obtained for each pixel along the line. Plots of pixel intensities across the yellow lines in Figures 6 and 7 are shown in the right panels. At least three independent transformations were performed for each plasmid, including appropriate controls each time.

## Acknowledgements

We acknowledge the ImagoSeine core facility of Institut Jacques Monod, member of France-BioImaging (ANR-10-INBS-04) and IBiSA, with the support of Labex “Who Am Iℍ, Inserm Plan Cancer, Region Ile-de-France and Fondation Bettencourt Schueller.

Computational resources were provided by the e-INFRA CZ project (ID:90254), supported by the Ministry of Education, Youth and Sports of the Czech Republic. This study was supported by the CNRS, France and grant ANR-20-CE13-0007 from the ANR, France to CLJ. Work in the Dacks lab is supported by grants from the Natural Sciences and Engineering Research Council of Canada (RES0043758 and RES0046091).

## Supplementary Figure Legends

**Supplementary Figure 1.**
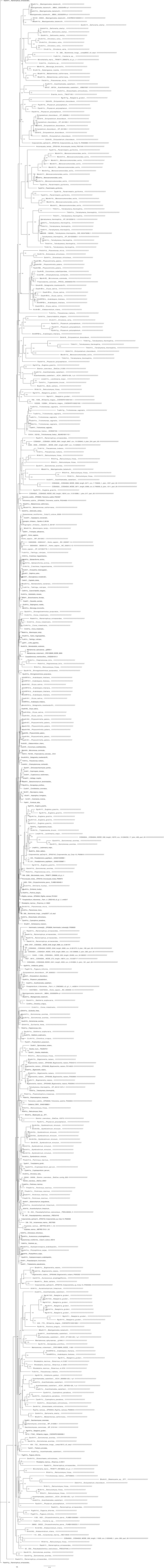
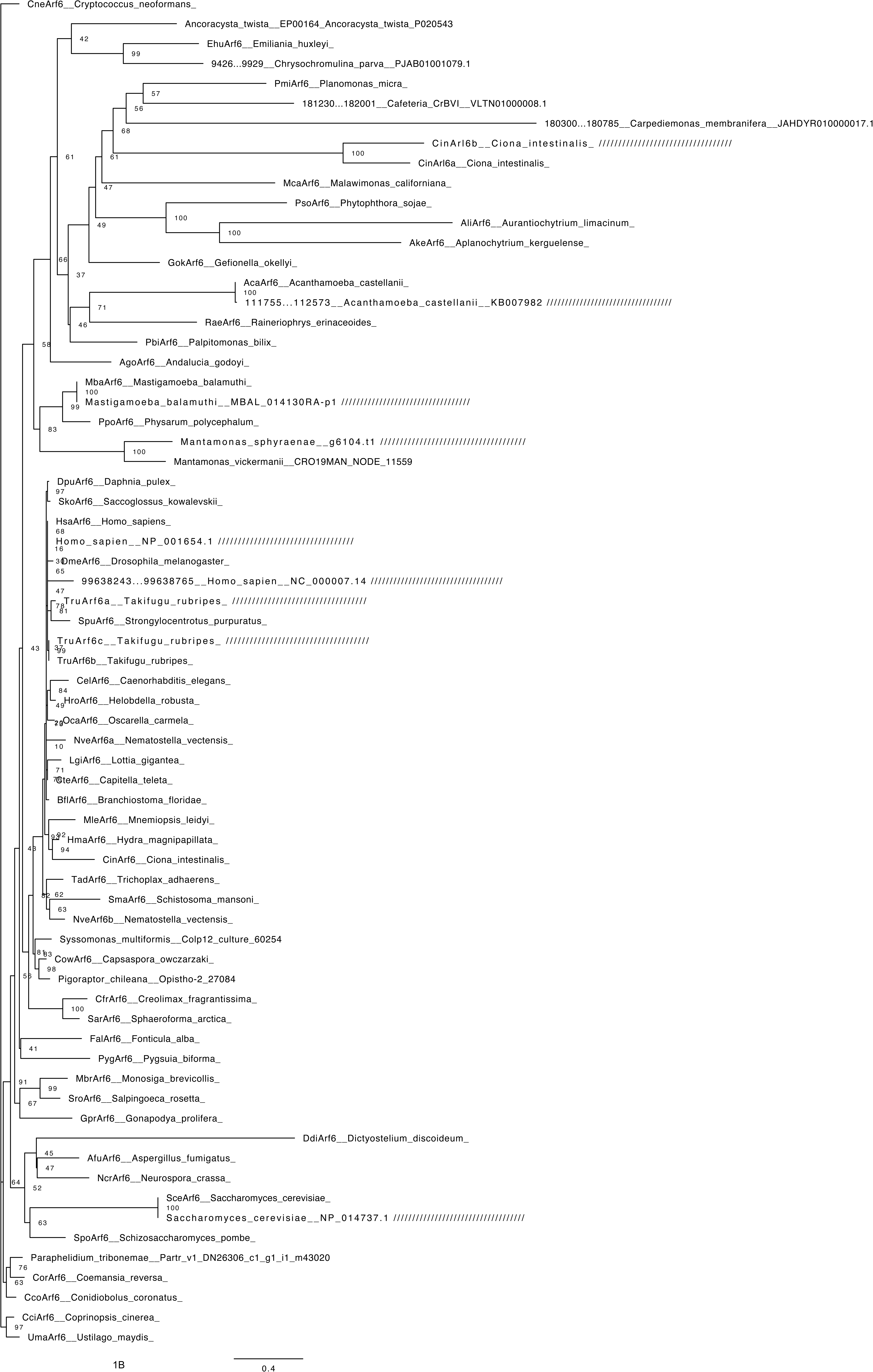
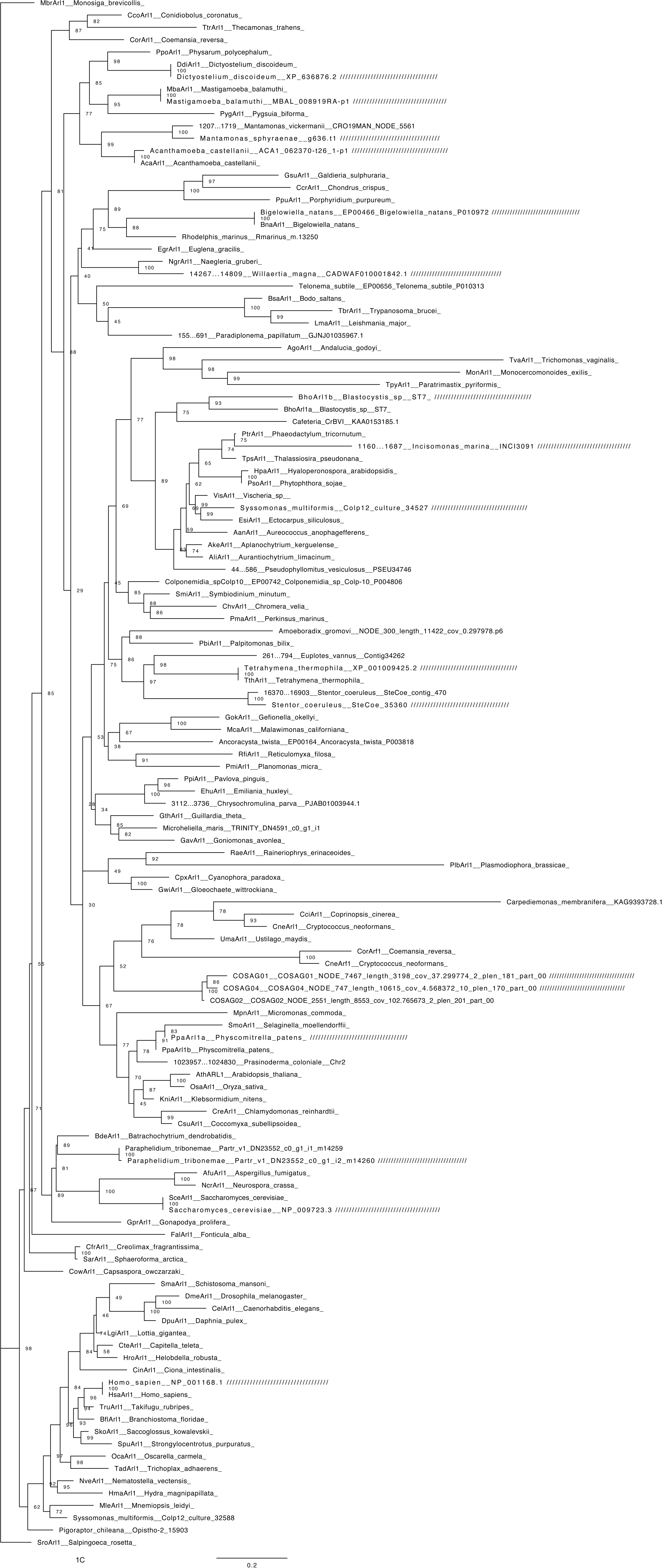
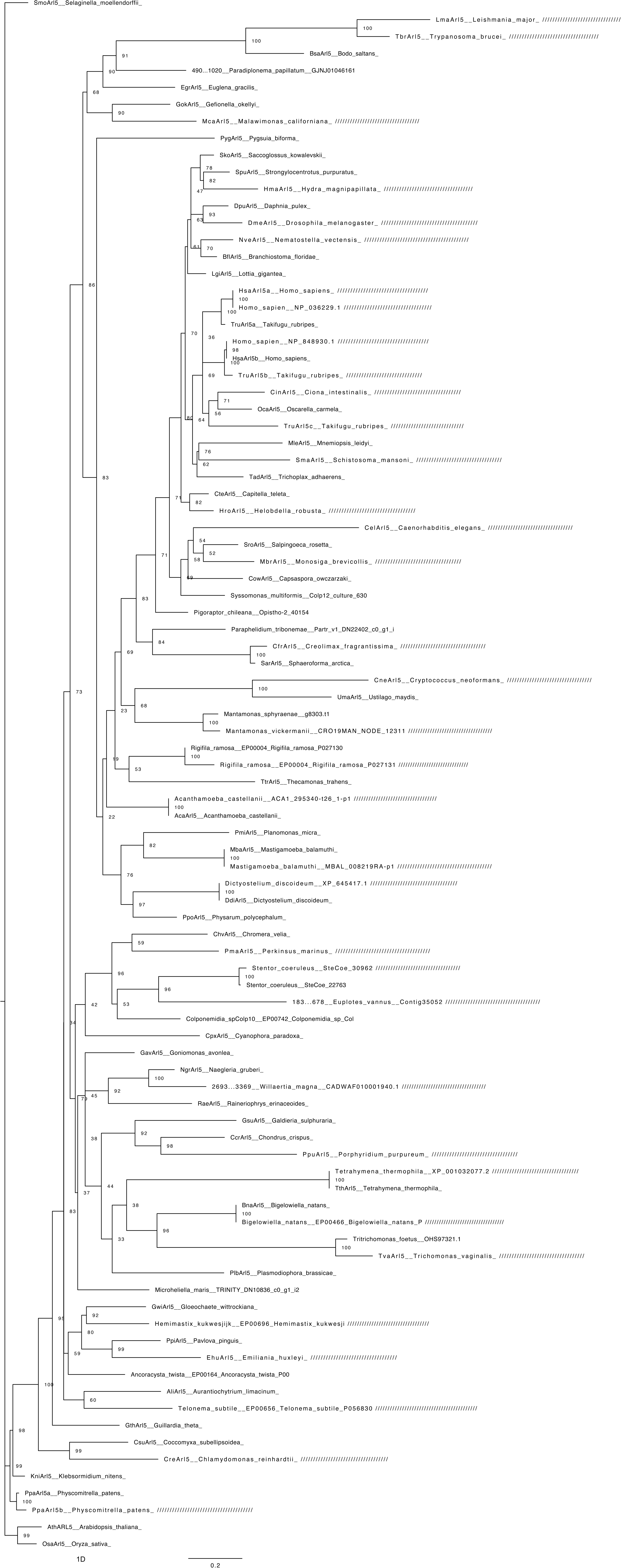
Phylogenetic analyses of Arf sub-families to identify least divergent paralogs. (A) Arf1. Model=Q.Insect+R8 Matrix=451 taxa, 171 positions. (B) Arf6 Model=Q.Insect+I+G4 Matrix=68, 170 positions. (C) Arl1. Model=Q.Yeast.R5 Matrix=126 taxa, 163 positions (D) Arl5. Model=Q.Yeast.R6. Matrix=92 taxa, 161 positions.

**Supplementary Figure 2.**
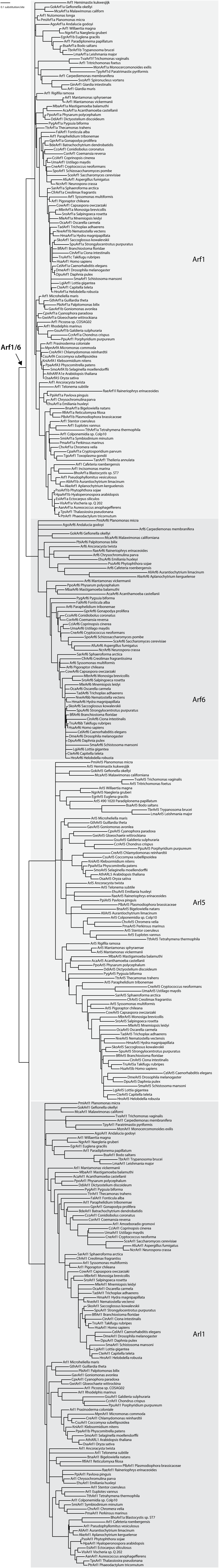
Constrained tree topology for ASR analyses. This tree was manually constrained based on the topology from Vargová et al. 2021 ^12^, assuming that Arl1/5 represents an outgroup root. Internal eukaryotic relationships within each clade are based on consensus eukaryotic relationships ^12^ and with a polytomy at the root, reflecting the knowledge at the time of analysis. Notably, the Carpediemonas ‘Arf6’ was included in the calculation yielding the sequences tested. However, recalculation without the sequence resulted in only a single AA residue change at a low probability site, and thus did not substantially change the inferred Arf1/6 ancestral sequence (Supplementary Table 2).

**Supplementary Figure 3.**
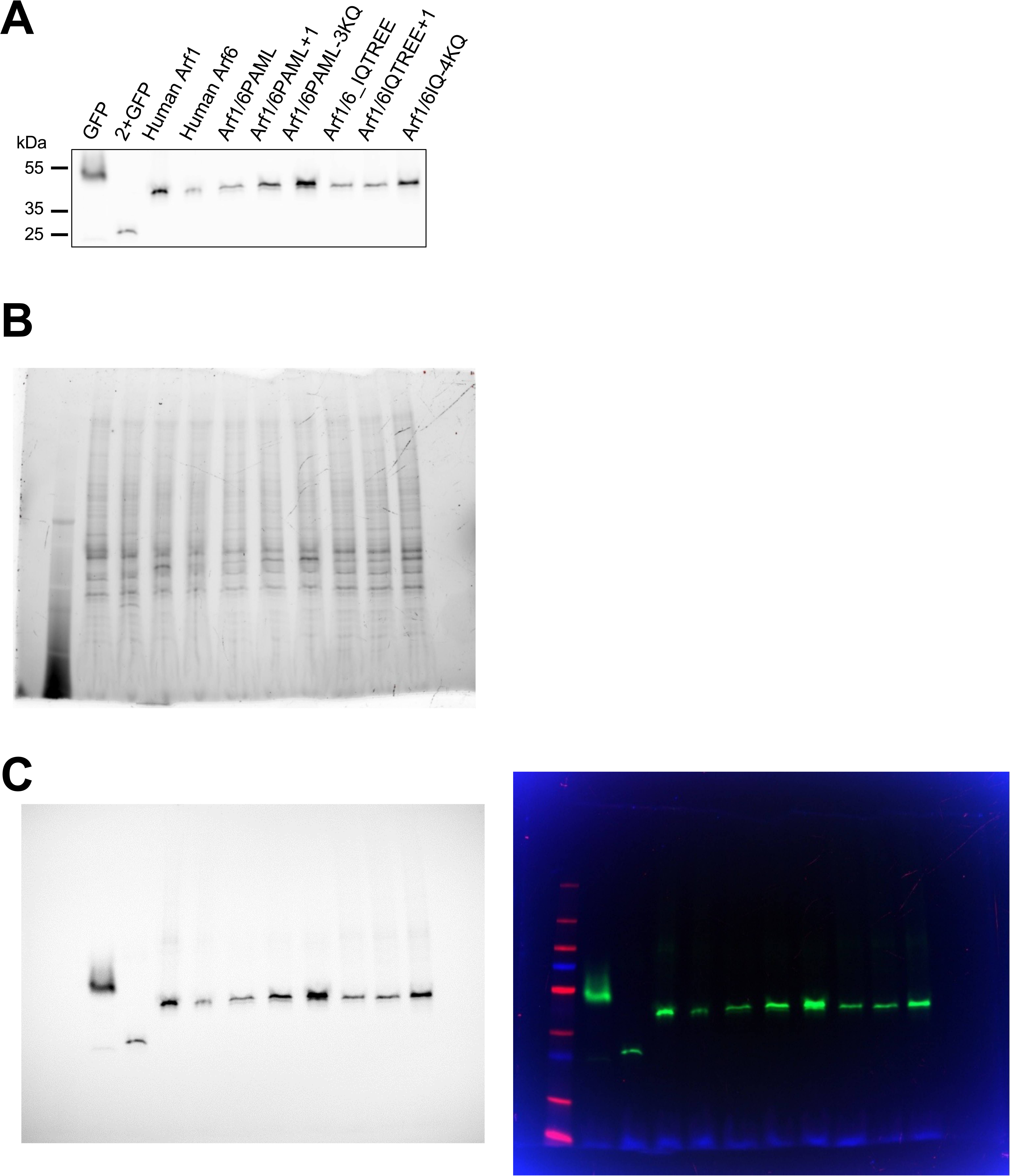
Expression levels of Arf1/6 predicted ancestral proteins and controls. MDCK cells were transfected with the plasmids encoding the indicated proteins fused to GFP (or GFP alone) (A), and samples prepared for GFP in gel fluorescence (see Methods) (A). As a loading control, total proteins were visualized in commercial pre-cast SDS–PAGE stain-free gels. (B). Uncropped version of blot in Part A is shown (C).

**Supplementary Figure 4.**
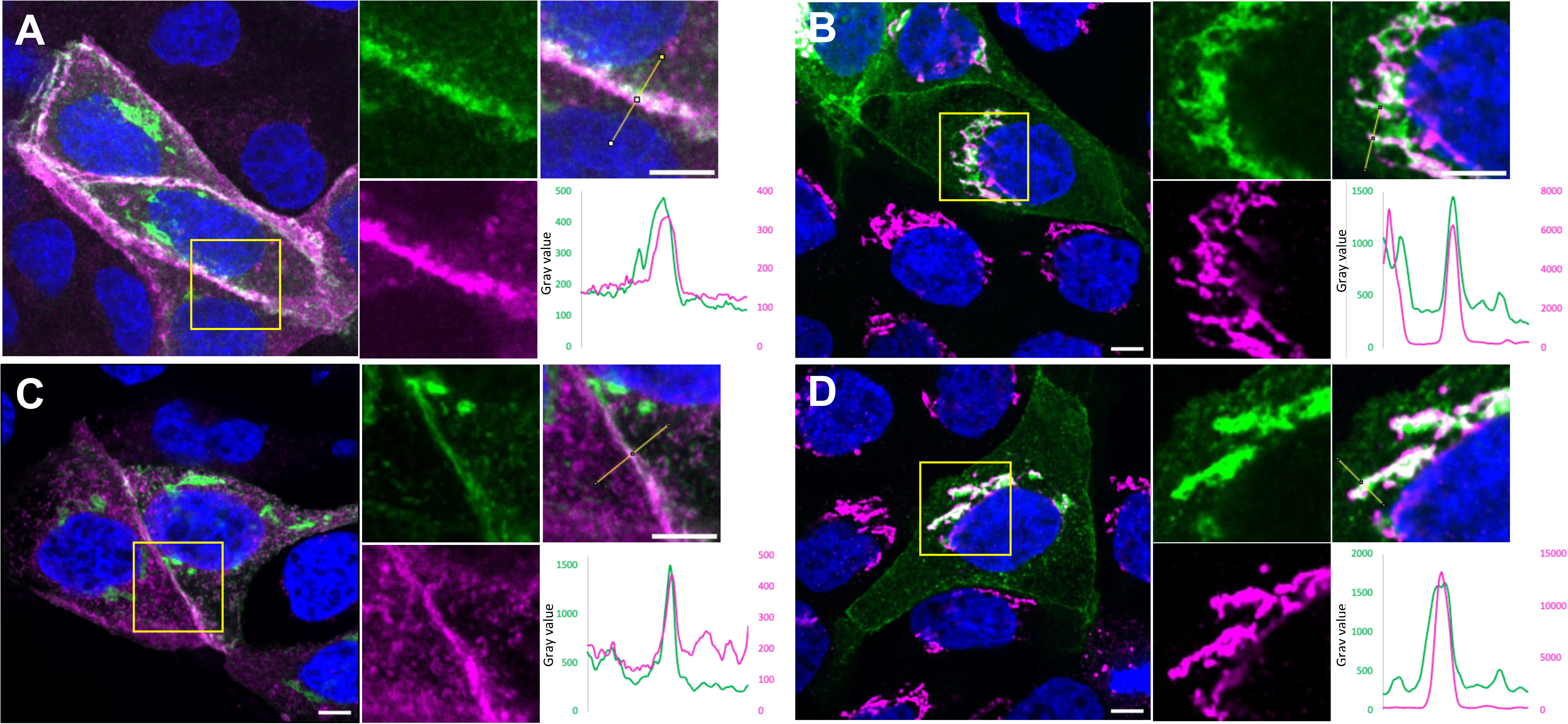
Localization of alternative Arf1/6 predicted ancestral proteins to the Golgi and PM in MDCK cells. MDCK cells were transfected with ARF1/6PAML+1-GFP (A,B) or ARF1/6IQ+1-GFP (C,D) and prepared for immunofluorescence analysis using Hoechst dye to visualize nuclei (blue). Cells were either co-transfected with the 8+mCherry PM probe (magenta) (A,C) or prepared for immunofluorescence analysis using antibodies against GM130, an early Golgi marker (magenta) (B,D). Representative merged images are shown on the left for each transformant, with GFP-tagged protein (green), nuclei (blue), and either PM (A,C) or Golgi (B,D) in magenta. Insets (yellow square) are shown on the right, with each channel shown separately (left inset panels). The yellow lines shown in the merged inset panels (upper right) are those used to generate plots of pixel intensities for the GFP-tagged proteins (green) and the organelle marker (magenta) (lower right) of each part (A-D). At least three independent transformations were performed for each mutant plasmid, always including the appropriate control plasmids. Scale bar, 5 μm.

**Supplementary Figure 5.**
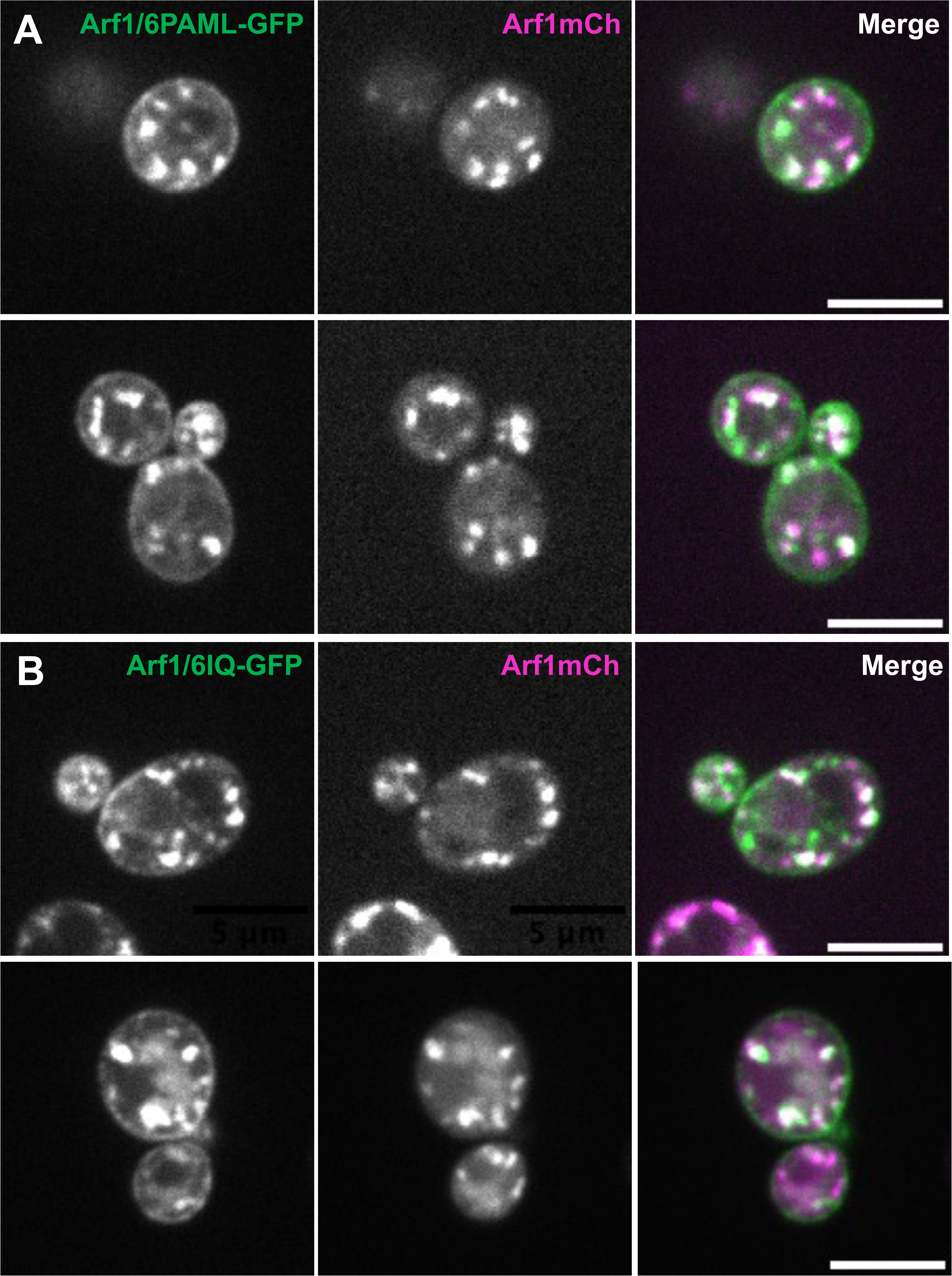
Localization of alternative Arf1/6 predicted ancestral proteins in yeast cells. Plasmids encoding Arf1/6PAML+1-GFP (A) and Arf1/6IQ+1-GFP (B) were co-transformed into *Saccharomyces cerevisiae* cells with a plasmid encoding Arf1-mCherry. Representative images for each GFP-tagged protein expressed (left panels), with Arf1-mCherry (middle panels) are shown. The merged images are shown in the right panels. At least three independent transformations were performed for each mutant plasmid, always including the WT control plasmid. Scale bar, 5 μm.

